# Bell-shaped dose response for a system with no IFFLs

**DOI:** 10.1101/2020.11.17.387605

**Authors:** Eduardo D. Sontag

## Abstract

It is well known that the presence of an incoherent feedforward loop (IFFL) in a network may give rise to a steady state non-monotonic dose response. This note shows that the converse implication does not hold. It gives an example of a three-dimensional system that has no IFFLs, yet its dose response is bell-shaped. It also studies under what conditions the result is true for two-dimensional systems, in the process recovering, in far more generality, a result given in the T-cell activation literature.

## 1 Introduction

For ± signed directed graphs, an “incoherent feedforward loop” (IFFL) is a pair of simple (not transversing any node twice) directed paths *P*_1_ and *P*_2_ from a node *N*_1_ to a node *N*_2_ between so that *P*_1_ and *P*_2_ have opposite net signs (product of signs along edges). For example, the 3-node signed graph in Figure 1 has an IFFL.

**Figure 1:**
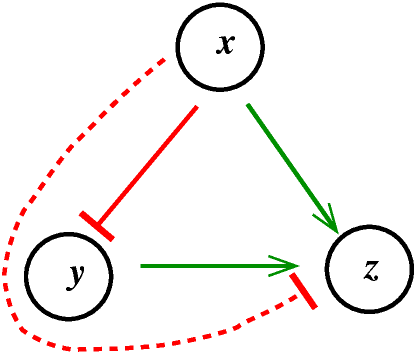
An example of an IFFL. The two solid green edges are positive, and the solid red edge is negative. The dashed path from node *x* to node *z* has a net negative sign, but the solid green path from node *x* to node *z* has a positive sign.

IFFLs are a ubiquitous motif in systems biology [1]. One associates a signed directed graph to a system by considering the Jacobian of the vector field. We assume here that any such Jacobian entry has a definite sign (≥ 0 or ≤ 0) throughput the state space. It is well-known that the presence of IFFLs may result in steady state dose responses which are non-monotonic; see e.g. [4] or the references in [7]. Negative feedbacks *or* IFFLs are necessary, as otherwise the theory of monotone systems implies that the dose response (“input to state characteristic”) will be monotone on input values [2, 3]. (In fact, for monotone systems, even transient responses at any given time also behave monotonically on input magnitude.)

It has been argued that IFFLs are also *necessary*. For example, in [5] the authors state that *“models without an incoherent feed-forward loop but with negative feedback… cannot produce a bell-shaped dose-response”* (main text as well as section “Negative feedback cannot produce a bell-shaped dose-response” in SI). This conclusion was based on a computational screen of 58,905 network architectures, as well as explicit examples in the SI. We show here that this converse is false, i.e. one may have non-monotonic dose-responses even in systems that include no IFFLs (but, of course, include negative feedback loops, as these are necessary if there are no IFFLs). Our counterexample is not included in the class of reaction formulas systems computationally explored in [5], thus explaining the discrepancy.

We also study under what conditions the result is true for two-dimensional systems, in the process recovering, in far more generality, a result given in [5] in the context of T-cell activation.

## 2 A counterexample for three-dimensional systems

Consider the following system:

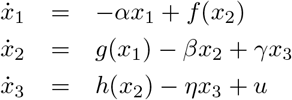

where the functions *f*, *g*, and *h* are decreasing and non-negative and *α, β, γ, η* are positive constants. The system evolves on states *x*_*i*_(*t*) ≥ 0, and the input *u* is taken as a constant in some interval *I* ⊆ [0, ∞). The output is *y* = *h*(*x*_1_, *x*_2_, *x*_3_) = *x*_3_. Note that the system is positive, meaning that when starting from nonnegative *x*_1_, *x*_2_, *x*_3_ the solutions remain nonnegative.

The Jacobian matrix (with respect to the variables *x*_*i*_), evaluated at any state, has the following sign structure:

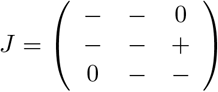

which we represent graphically as in Fig. 2.

**Figure 2:**
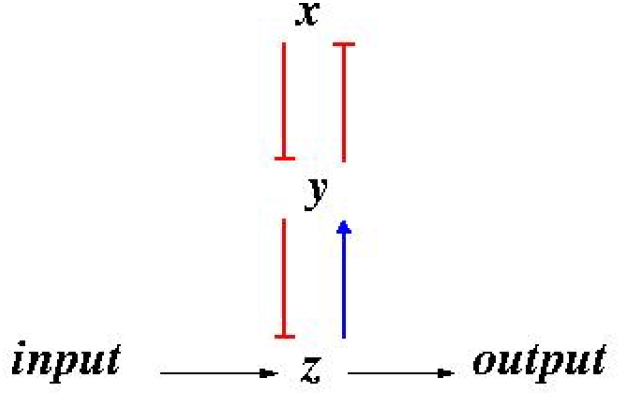
Signed digraph representing the three-dimensional example. (Using *x, y, z* instead of *x*_*i*_’s.)

Clearly, there and no IFFLs in our network, because there is only one directed path between any two pairs of nodes.

In general, suppose that for each constant input value *u* ∈ *I* there is an asymptotically stable steady state *X*_*u*_. The output at this steady state is *y* = *h*(*X*_*u*_), and the mapping *u* ↦ *h*(*X*_*u*_) is defined as the *steady state dose response* of the system. We will now provide functions *f*, *g*, and *h* for which the dose response for our system is non-monotonic (see right panel in Figure 3).

**Figure 3:**
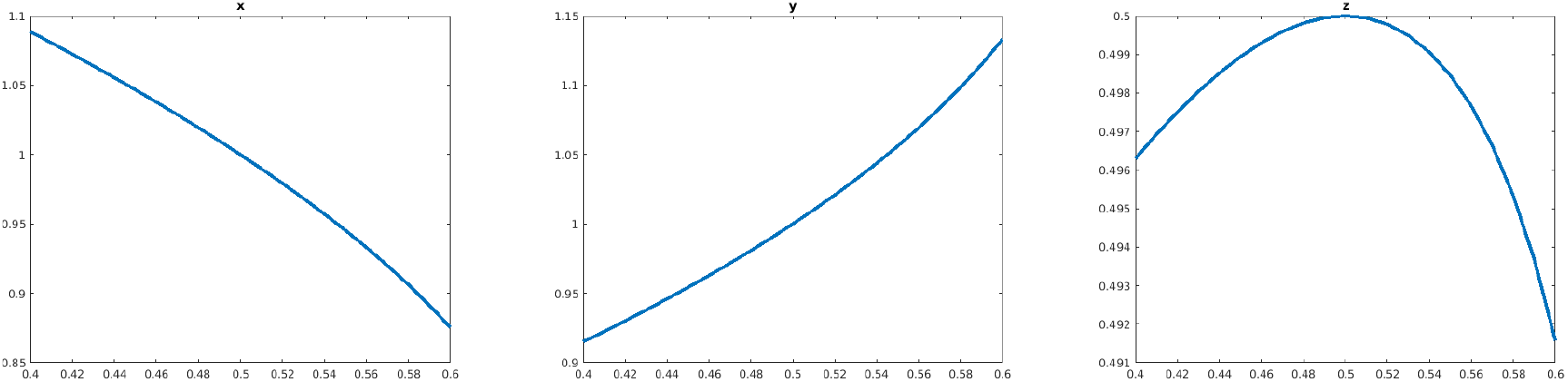
Plots of steady states *x*_1_, *x*_2_, *x*_3_ (labeled *x, y, z* in figure) as functions of constant input. Note the bell-shape of the output *x*_3_.

We define the functions *f* and *g* as follows:

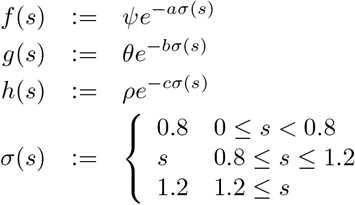

and the positive parameters are as follows:

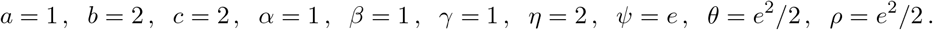

That is, the system is:

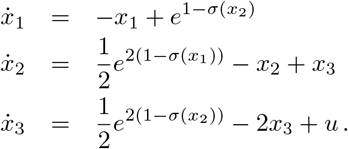

We study equilibria for constant inputs in the interval

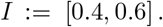

At the midpoint of this interval, *u*_0_ = 0.5, a steady state is

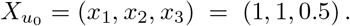

We now find equilibria for any given constant input *u*. From 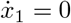 we have that

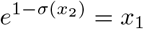

and then solve 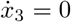 for *x*_3_, using that 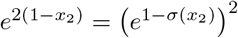, and obtain:

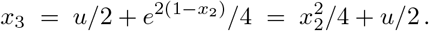

Finally, we set 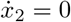, substituting the above *x*_3_, to obtain an equation relating *x*_1_, *x*_2_, and *u*:

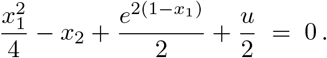

This equation is equivalent to

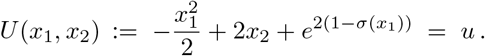

Now we will show that, for each *u* ∈ *I*, there is a unique solution (*x*_1_, *x*_2_) for *U* (*x*_1_, *x*_2_) = *u*, and, moreover, this solution has *x*_1_ ∈ [0.8, 1.2] and *x*_2_ ∈ [0.8, 1.2], the non-saturation regime, case (1A) below. We do this by analyzing various cases.

**Case 1:** *x*_2_ ∈ [0.8, 1.2].

(1A) Consider first the case *x*_1_ ∈ [0.8, 1.2]. Thus *σ*(*x*_*i*_) = *x*_*i*_ for *i* = 1, 2. Now *x*_2_ = 1 − ln *x*_1_, so we substitute and write *U* as a function of *x*_1_ alone:

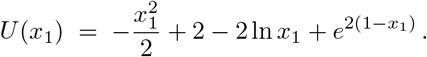

This is a strictly decreasing function of *x*_1_ on the interval [0.8, 1.2] and its values range from 0.6345 at *x*_1_ = 0.8 to 0.2450 at *x*_1_ = 1.2. Thus, the interval *I* = [0.4, 0.6] is included in the range of *U*, and a unique solution of *U* (*x*_1_) = *u* exists. From this, we obtain the unique values *x*_2_ = 1 − ln *x*_1_ and then 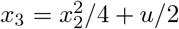.

(1B) Assume now *x*_1_ < 0.8, so *σ*(*x*_1_) = 0.8. Now

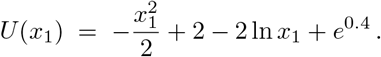

This is a strictly decreasing function of *x*_1_ on the interval [0, 0.8] and its values range from +∞ at *x*_1_ = 0 to 0.6345 at *x*_1_ = 0.8. Therefore *I* = [0.4, 0.6] does not intersect the range. Thus there are no solutions in case (1B).

(1C) Assume now *x*_1_ > 1.2, so *σ*(*x*_1_) = 1.2. Now

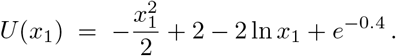

This is a strictly decreasing function of *x*_1_ on the interval [1.2, ∞) and its values range from −0.5765 at *x*_1_ = 1.2 to −∞ at *x*_1_ = +∞. Therefore *I* = [0.4, 0.6] does not intersect the range. Thus there are no solutions in case (1C).

**Case 2:** *x*_2_ ∈ [0, 0.8], so *σ*(*x*_2_) = 0.8. Now 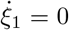 means that 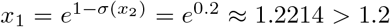 and thus the only case for *x*_1_ to consider is *x*_1_ ≈ 1.2214, *σ*(*x*_1_) = 1.2. So

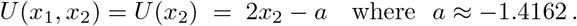

This is an increasing function of *x*_2_ on the interval [0, 0.8], and its values range from ≈ −1.4162 at *x*_2_ = 0 to 0.1838 at *x*_2_ = 0.8. Therefore *I* = [0.4, 0.6] does not intersect the range. Thus there are no solutions in case (2).

**Case 3:** *x*_2_ > 1.2, so *σ*(*x*_2_) = 1.2. Now 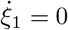 means that 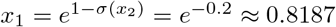 and thus the only case for *x*_1_ to consider is *x*_1_ ≈ 0.8187, *σ*(*x*_1_) = *x*_1_ ≈ 0.8187. So

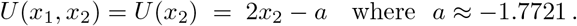

This is an increasing function of *x*_2_ on the interval [1.2, +∞), with a minimum value of ≈ 0.6279 at *x*_2_ = 1.2. Therefore *I* = [0.4, 0.6] does not intersect the range. Thus there are no solutions in case (3).

We conclude that steady states are unique, and occur only in case (1A), when neither variable is saturated.

For case (1A), then, we solve numerically

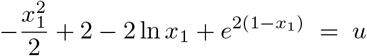

to get *x*_1_ = *x*_1_(*u*) for each constant input *u* in the interval *I* and back substitute to obtain the remaining variables, obtaining a steady state *X*_*u*_, Figure 3, We verify that the dose response (third coordinate) is indeed not monotone. See Figure 3.

The maximal real part of the eigenvalues of the Jacobian matrix along these steady states is always negative, showing asymptotic stability, Figure 4.

**Figure 4:**
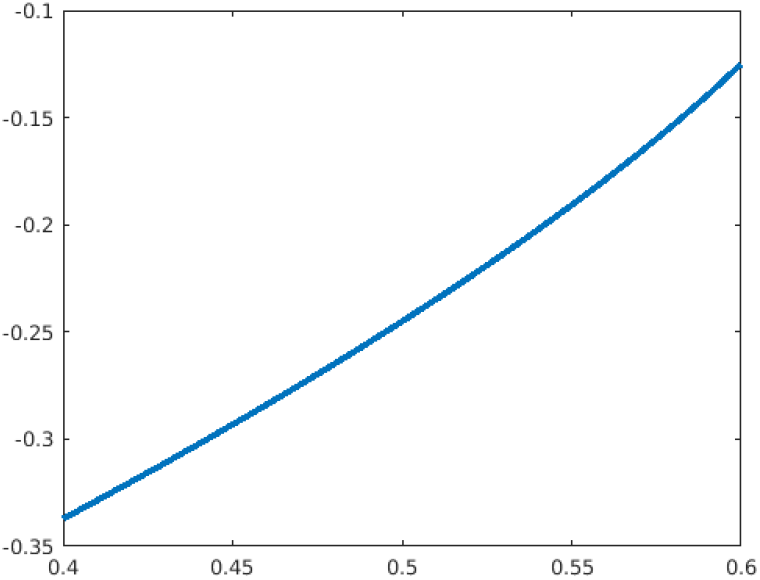
Maximum real part of eigenvalues, as a function of *u*, showing stability.

Simulations from random initial states indicate that the state *X*_*u*_, *u* ∈ *I* is globally asymptotically stable.

### Remark

A small technical issue is that the saturation *σ*, while a (globally) Lipschitz function (insuring unique and everywhere defined solutions of the ODE) is not differentiable at those states for which either *x*_1_ or *x*_2_ equals exactly 0.8 or 1.2, which happens only a set of measure zero. Thus the Jacobian is not well-defined at every state in the classical sense. However, it is well-defined in the sense of nonsmooth analysis, and in any event the signs of interactions reflect the increasing or decreasing influence of each variable on each other variable, and this can be defined with no need to take derivatives. The saturation function *σ* could be replaced by an approximating differentiable function, but use of *σ* makes calculations somewhat simpler.

This example was derived as follows.

We first ignore the saturations, and consider the Jacobian (with respect to the *x* variables):

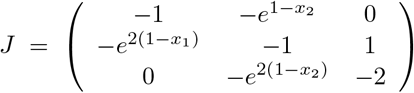

which has characteristic polynomial *P* (*λ*) = *λ*^3^ + *a*_2_*λ*^2^ + *a*_3_*λ* + *a*_4_ with

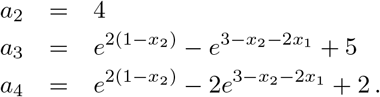

the Routh-Hurwitz criterion requires that *a*_2_, *a*_3_, *a*_4_ > 0 and Δ ≔ *a*_2_*a*_3_ − *a*_4_ > 0 for stability. In particular, when *x*_1_ = 1 and *x*_2_ = 1 (as in the equilibrium *X*_*u*_ corresponding to *u* = 0.5), *a*_2_ = 4, *a*_3_ = 5, and *a*_4_ = 1, so these properties are satisfied here, and that this equilibrium is stable.

It follows that, in particular, the Jacobian is nonsingular at this equilibrium, so that, by the implicit function theorem, we know that there is a local curve *X*_*u*_ of equilibria for each *u* near *u* = 0.5. We then guessed a range where stability will be preserved, by checking the Routh-Hurwitz criterion as a function of (*x*_1_, *x*_2_). This suggested the range to be used, and the eigenvalue plot confirmed the results. The saturation function was added in order to provide unique equilibria and thus the possibility of global stability; without it, we found that there is bistability.

The fact that the dose response is bell-shaped is suggested by a perturbation argument, with no need to plot numerically. In terms of the derivatives 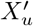 of *X*_*u*_ with respect to *u*, we have an implicit equation

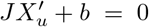

where *b* = col(0, 0, 1). Thus the derivative of the output *y* = *x*_3_ is, as a function of *u*:

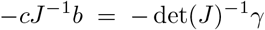

where *γ* is the (3, 3) entry of the adjugate matrix, which is the principal 2 × 2 minor of *J*, namely zero. Thus we have a point at which the output may have a local minimum or maximum. One could check the second derivative to see that this is strict, but at this point a simulation completes the proof.

### Remark

In any dimension, for systems 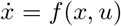 with output *y* = *h*(*x*), one may similarly check by a perturbation argument if the determinant of the Jacobian of *f* and a suitable minor do not change sign along the state space. A sufficient condition for this can be given in terms of regular interval matrices [6]. We omit the easy details.

### Remark

The bell-shaped dose response in Fig 3 (right panel) spans only a small range of output values. One could rescale this example to have a more dramatic effect. There are at least two ways to change this while still having the same result (no IFFL):

- We can add a new variable, let us say *x*_4_, with an equation 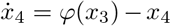, where now the output is *x*_4_, and the function *φ* is strictly increasing. This system will still have no IFFLs. Now the dose-response is *φ*(*h*(*X*_*u*_)), where *h*(*X*_*u*_) was the previous dose response. We can take a Hill function for *φ*. rendering the new dose response initially almost 0, changing to 1, and finally back to near zero.
- A less elegant alternative, if we want to insist on three-dimensional example, is to make a change of variables *x*_3_ = *φ*(*x*_3_), with the same function *φ* as above. Now the equation for 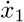 will have a term *φ*^−1^(*x*_3_), which is still increasing on *x*_3_, and the equation for 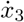 will have the term *φ*′(*φ*^−1^(*x*_3_)) multiplying the previous left hand side, so all Jacobians still have the same sign.

## 3 The case of two-dimensional systems

Consider this diagram for a system, containing one negative feedback loop (and no IFFL):

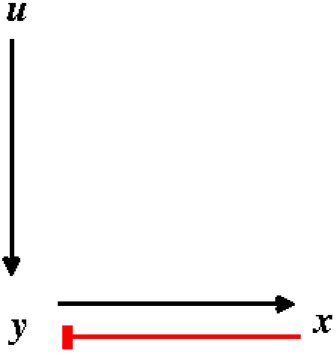

Here *u* is the input, *y* the output, and *x* the intermediate state that provides the negative feedback loop. We model this as

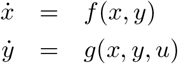

on *x*(*t*) ≥ 0, *y*(*t*) ≥ 0 (with *u* ≥0 seen as a constant in this problem), and we impose these conditions (subscripts indicate partial derivatives):

1. *f*_*y*_(*x, y*) > 0 for all *x, y* (arrow from *y* to *x* is activating),
2. *g*_*x*_(*x, y, u*) ≤ 0 for all *x, y, u* (arrow from *x* to *y* is repressing),
3. *g*_*u*_(*x, y, u*) ≥ 0 for all *x, y, u* (arrow from *u* to *y* is activating),
4. *f* (0, *y*) ≥ 0, *g*(*x,* 0, *u*) ≥ 0 for all *x, y, u* (invariance of nonnegative orthant).

In addition, we impose these constraints to make the problem non-trivial:

5. for each non-negative constant input *u*, there is a unique (non-negative) steady state, denoted as (*x*_*u*_, *y*_*u*_)
6. (*x*_*u*_, *y*_*u*_) depends smoothly on *u* (this is done for purely technical reasons, to be able to compute the sensitivity on *u*),
7. this steady state is exponentially asymptotically stable.

We call *y*_*u*_ as a function of *u* the *dose response*.

We will show:

- The *dose response need not be monotone* (it may be bell-shaped).
- However, *if we impose the additional requirement* that *f*_*x*_(*x, y*) ≤ 0 for all *x, y*, then the dose response *is monotone*.

This last requirement says, in effect, that *x* can degrade but there is no auto-catalysis on *x*.

We prove the second statement and give an example for the first one.

The proof of is immediate by the arguments given above - now the adjugate matrix is just a single element that is always negative. We prove it in detail.

We compute the Jacobian:

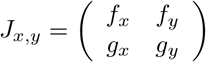

The assumed stability condition means that the trace is negative and the determinant is positive (at the equilibrium (*x*_*u*_, *y*_*u*_), for each *u*):

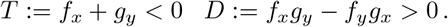

(Observe that, if we assume that both *f*_*x*_ ≤ 0 and *g*_*y*_ < 0, then we automatically have *T* ≤ 0 and *D* ≥0, but we do not need these additional assumptions, since we are assuming stability.)

The steady states satisfy

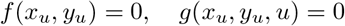

at all *u*, and are differentiable in *u*, so we differentiate:

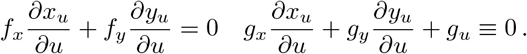

Thus

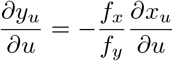

and

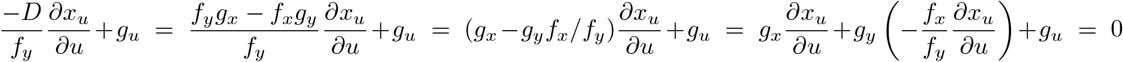

which gives

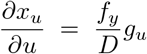

and thus substituting back in the formula for 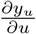:

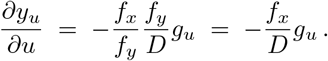

Therefore, if we also have *f*_*x*_ ≤ 0 then 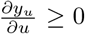 and we conclude that *y*_*u*_ is increasing on *u*. (Strictly if *f*_*x*_ < 0 and *g*_*u*_ > 0.)

This ends the proof.

### Remark

Note that not all the properties 1-7 are used in the proof. Also, the same result holds if we have an invariant set 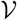 for the solutions of the differential equations and we are only interested in solutions that remain in 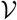. Of course, the conditions are then imposed on all 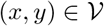.

### Remark

In the paper [5] the authors prove the same result for the particular system evolving on [0, 1]^2^ in which:

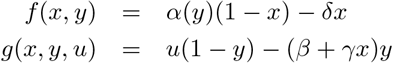

and

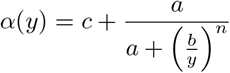

for certain (positive) parameters *a, b, c*. (For simplicity, we redefined the input in that paper, to absorb an additive and a multiplicative constant.) All conditions for our result are satisfied. (Note that *f*_*x*_ = *α*(*y*) *δ* < 0.) The existence of a unique steady state follows from the fact that one can solve the first equation to obtain *x* = *θ*(*y*) ≔ *α*(*y*)/(*δ* + *α*(*y*)) and when substituted into *u*(1 − *y*) = (*β* + *γθ*(*y*))*y* we have that the left hand side decreases from *u* to zero and the right had side increases from zero, so there is a unique equilibrium (and it is smoothly dependent on *u* by the implicit function theorem).

This construction also hints at the counterexample. We need to find *f* and *g* so that: 1-7 hold and *g*_*x*_ changes sign along the curve (*x*_*u*_, *y*_*u*_).

In order to obtain such a counterexample, we will take special forms for *f* and *g*:

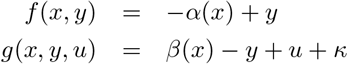

where *κ* > 0 is a constant (this could be though of either as constitutive production of *y* or as a lower bound on a redefined input *κ* + *u*) and the smooth functions *α* and *β* are defined below.

We need the following properties:

1. *f*_*y*_(*x, y*) > 0 (satisfied because *f*_*y*_ = 1 > 0)
2. *g*_*x*_(*x, y, u*) ≤ 0, i.e. *α*′ ≥ 0 (here prime is derivative w.r.t. *x*)
3. *g*_*u*_(*x, y, u*) ≥ 0 (satisfied because *g*_*u*_ = 1)
4. *f* (0, *y*) ≥ 0, *g*(*x,* 0, *u*) ≥ 0: we’ll take *α*(0) = 0 and *β*(*x*) ≥ 0

and additional properties as follows.

The steady state for constant input *u* solves these two equations:

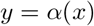

and

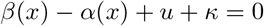

or equivalently

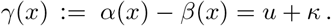

We will require that, for some *δ* > 0, *γ*(*δ*) = *κ* and *γ*(*x*) > 0, *γ*′(*x*) ≥ 0 for all *x* ∈ [*δ,* ∞). By the implicit mapping theorem, this says that there is a unique (and differentiable) solution *x* ∈ [*δ,* ∞) of *γ*(*x*) = *u* + *κ*, for each *u* ≥ *κ*, and this solution depends smoothly on *u*. This will be *x*_*u*_. We will also require *α*(*x*) ≥ 0 for all *x* ∈ [*δ,* ∞], which then gives *y*_*u*_ = *α*(*x*_*u*_). Thus, there is a unique steady state for each *u*, and (*x*_*u*_, *y*_*u*_) depends smoothly on *y*, guaranteeing gives properties 5 and 6. There remains to assure stability: *D* > 0 and *T* < 0 at all equilibria, Since *f*_*x*_ = −*α*, *f*_*y*_ = 1, *g*_*x*_ = *β*′, *g*_*y*_ = −1:

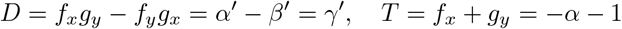

we need to require *α*(*x*) > −1 for all *x* ∈ [*δ,* ∞) (we already required *γ*′ > 0). To summarize, we need:

- smooth functions *α, β*: [0, ∞) → ℝ,
- *α*(0) = 0,
- *β*(*x*) ≥ 0 for all *x* ∈ [0, ∞),
- numbers *δ* > 0, *κ* > 0 so that *γ* ≔ *α* − *β* has *γ*(*δ*) = *κ*,
- *γ*′(*x*) > 0 for all *x* ∈ [*δ,* ∞) (this implies that *γ*(*x*) ≥ *γ*(*δ*) > 0)
- *α*(*x*) > −1 for all *x* ∈ [*δ,* ∞).

Our objective is to get a sign change in 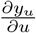, so we need that *f*_*x*_ changes sign, i.e. *α*′ should admit both a positive and a negative sign on *x* ∈ *x* ∈ [*δ,* ∞).

There are many possible choices for such *α* and *β*. We pick a rather artificial one merely to show the existence of such functions, not for any practical significance. We will pick

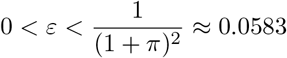

and any

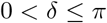

and these functions:

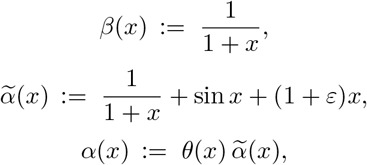

where *θ* is a smooth function which is 0 at *x* = 0 (thus assuring *α*(0) = 0) and *θ*(*x*) ≡ 1 for *x* ≥ *δ*. Observe that this definition assures that 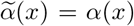 for all *x* ≥ *δ*. Such functions *θ* can be obtained by an abstract partition of unity argument. However, the same argument can be adapted to work with a Hill function *θ*(*x*) = *x*^*n*^/(*K*^*n*^ + *x*^*n*^) with 0 ≪ *K* ≪ 1 and *n* ≫ 1.

Note that 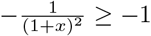 and cos *x* ≥ −1, so the function 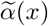 has derivative

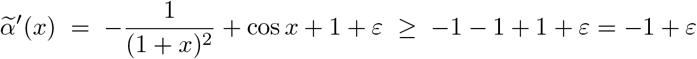

which shows that 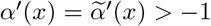 for *x* ≥ *δ*.

Let 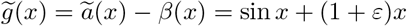, and that 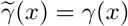 for all *x* ≥ *δ*. Observe that

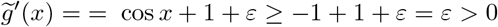

for all *x* and that *γ*(0) = 0, so it follows that 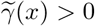 for all *x* (and thus in particular for *x* ≥ *δ*).

When *x* = π, 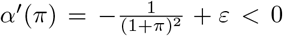 because of the choice of *ε* On the other hand, for *x* = 2π, 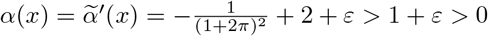. This shows that *α*′ changes sign.

We now-define

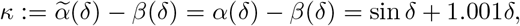

so that *γ*(*κ*) = *δ* is satisfied.

For example, if we take *ε* = 0.001 and *δ* = *π*/2, *κ* = 1 + 1.001*π*/2.

As a numerical example, with *ε* = 0.001 and

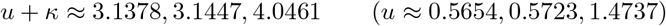

which were obtained by evaluating *γ*(*x*) at the respective points below:

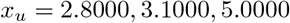

and the does response is not monotone, since we have these respective values for *y*:

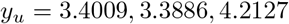

(we built up this example by starting from the fact that 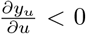 in an interval around an *u* for which *x*_*u*_ = *π*, since the sign of 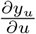 is the sign of *α*′ at that point). See Fig. 5 for part of the plot of *y*_*u*_ vs *u*.

**Figure 5:**
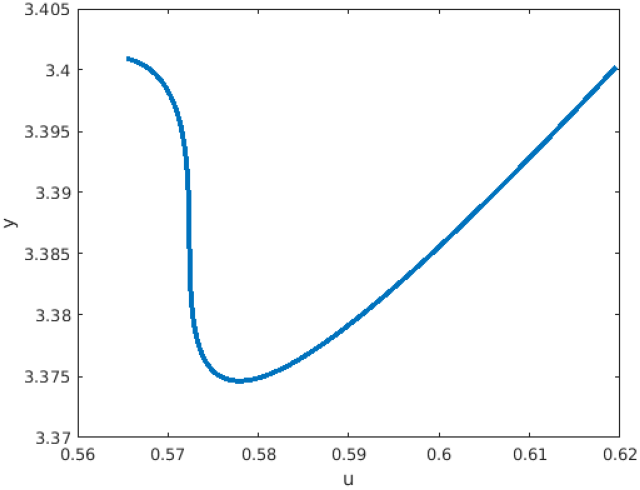
Non-monotone dose-response for two-dimensional example

## References

[1] U. Alon. An Introduction to Systems Biology: Design Principles of Biological Circuits. Chapman & Hall, 2006.

[2] D. Angeli and E.D. Sontag. Monotone control systems. IEEE Trans. Automat. Control, 48(10):1684–1698, 2003. Errata are here: http://sontaglab.org/FTPDIR/angeli-sontag-monotone-TAC03-typos.txt.

[3] D. Angeli and E.D. Sontag. Behavior of responses of monotone and sign-definite systems. In K. Hüper and Jochen Trumpf, editors, Mathematical System Theory - Festschrift in Honor of Uwe Helmke on the Occasion of his Sixtieth Birthday, pages 51–64. CreateSpace, 2013.

[4] D. Kim, Y. K. Kwon, and K. H. Cho. The biphasic behavior of incoherent feed-forward loops in biomolecular regulatory networks. Bioessays, 30:1204–1211, Nov 2008.

[5] M. Lever, H.S. Lim, P. Kruger, J. Nguyen, N. Trendel, E. Abu-Shah, P. K. Maini, P. A. van der Merwe, and O. Dushek. Architecture of a minimal signaling pathway explains the T-cell response to a 1 million-fold variation in antigen affinity and dose. Proc. National Academy of Sciences USA, 113(43):E6630–E6638, 2016.

[6] G. Rex and J. Rohn. A note on checking regularity of interval matrices. Linear and Multilinear Algebra, 39(3):259–262, 1995.

[7] E.D. Sontag. Remarks on feedforward circuits, adaptation, and pulse memory. IET Systems Biology, 4:39–51, 2010.

